# Xenomai-based multiple-process system, for real-time data acquisition and graphical display control

**DOI:** 10.1101/191973

**Authors:** Hadrien Caron, Pierre Pouget

## Abstract

To elicit complex and rich graphical displays, and record neuronal phenomena ofinterest while all simultaneously being capable to interact in a closed-loop with external devises is a challenging task to all neurophysiologists. To facilitate this process, we have developed an Open-Source software system using a single computer running a well established Linux architecture (Ubuntu) associated to a kernel duo providing hard real-time support (Xenomai). We show that a single computer using our API is capable, for any tasks that require OpenGL displaying, to acheive millisecond accuracy programmed events. In this report, we describe the design of our system, benchmark and its performance in a real-world setting, and describe some key features.

## 1. INTRODUCTION

Eliciting graphical displays, while simultaneously recording neuronal phenomena ofinterest and interact in a closed-loop with some external devises is a challenging task to realize in a standertized commonly used computer. One of the first well-known systems to accomplish this task was the complex UNIX-based real-time Tet() application developed for oculomotor experiments by Hays et al. (1982). As new applications have continued to emerge, however, many have forsaken true (or hard) real-time control and response in favor of simplicity, extended functionality, alternative operating systems, andr user-friendly design. Those systems that have maintained real-time support (e.g., TEMPO, ePrime, Matlab Psychtoolbox) are often costly, proprietary, and require additional code in order to define specific experiments. Finally, other systems utilizing the proven LabVIEW development architecture (e.g., Kodosky and Dye, 1989; Kullmann et al., 2004; Poindessault et al., 1995; Pruehsner et al., 2003; Sakatani and Isa, 2004;Gandhi and Bonadonna, 2004) was developed and optimized. However while these systerffeoopen access configuration, displaying are largely limited due to an LED board design necessary to a system refresh rate of 1 kHz and lacking monitor driver’s development. By creating an open source system that has minimal programming overhead and allow OpenGL graphical developpement, we hoped to allow users to focus on the essential features of experimental design and the basic elements of behavioral control and monitoring rather than on the often arcane details of the video presentation and data acquisition hardware. Our major goals were:

- To allow behavioral control with high temporal precision in free open source ressources.
- To allow straightforward scripting of behavioral tasks using OpenGL and-C++ syntax and functions.
- To interface transparently with data acquisition hardware for input / output functions, such as analog continuous signal, joystick and button-press acquisition, reward delivery, digital event marker output, as well as analog and TTL output to drive stimulators and injectors.
- To allow the full reconstruction of task events from the behavioral data file by including complete descriptions of behavioral performance, the event markers and their text labels, the task structure, and the actual stimulus images used; as a demonstration of this goal, to allow the replaying of any given trial from the behavioral data file alone.
- To provide the experimenter with an information rich display of behavioral performance and to reflect task events in real-time to aid the assessment of on-going behavior.

Rexeno is a software we developed that seeks to bind all these characteristics into a single piece of program, using a single computer, taking advantage of the best Open Source ressources available. The main charactestic of this program is that it will be able to interract in real-time with every ressources connected to it (Screens, A/D cards, Hard Drive, etc…).

Our tested system was composed of a Dell Computer with an Intel Duo processor (model E5405) running at 2.00 GHz and containing 2048MB of RAM (Dell Inc., Round Rock, TX). The graphics hardware in this machine consisted of an nVidia ENGT250 silent with 1024MB of video RAM. Output from this dual-headed graphics card was split to two subject displays running in full-screen mode at pixel resolutions of 1024 768. The displays were standard cathode-ray tubes mea-suring 19 inches in the diagonal, also from Dell. The refresh rate for the tests reported here was 60 Hz and the experimenters display window was set to update every 16.6 ms during behavioral monitoring to allow near-real-time (< 1*msec*) observation of the subjects performance.

To assess the performance of our software, we first collected data from a photodiode coupled with our neurophysiological recording system (Plexon Inc, TX, USA). Thus, we analyzed data simple from the on-going training of two rhesus monkeys (macaca mulatta, male, respectivelly 10.5 and 16 Kg) in a saccade inhbitory task.

For the animal testing and in order to allow eye-tracking, head fixation was achieved using a titanium head-post system (Crist instrument, Hagerstown, MD, USA). Visual fixation was required for a total of 3s (about 1 s of initial fixation followed by a 1-1.500 ms target presentation. An inter-trial-interval of 1 to 2 s was used. Data from three consecutive months of training (one session each day) were collected and confirmed to yield nearly identical results, so one of these sessions was chosen arbitrarily for presentation below. This session consisted of 1500 trials over 2 hours. At all times, the animal was handled in accord with EU guidelines and those of the ICM Animal Care and Use Committee. Analog X and Y position signals conveying behavioral output con-sisted of an optical eye-tracking system (Eyelink 2k, SR Research Ltd., Mississauga, Ontario, Canada) running at 1000 Hz. For a more straightforward demonstration, a schematic diagram, the code (timing script) and the conditions file for a simpler task (a standard pro-saccade task) is shown in Figure **??**. The object of this paper is to explain how we designed and tested our Real Time System, then we will detail how to obtain and use it. Usability is a critical issue for us because we needed powerful tools designed by others in order to build Rexeno, and we hope it will also be adapted by others in order to help them in their respective tasks.

## 2. Material and Methods

### 2.1. Existing alternatives

One millisecond is a relatively course (small ?) unit of measure by electronic standards. However such temporal precision on a non-hard-real-time system, running on most popular operating systems (Windows, Mac, Unix), has no guarantees, because the predictability of software events is limited by the design of the operating system (OS). Specifically, even those processes designated as having a real-time priority can be pre-empted by both kernel-level events and by interrupt requests, as well as by other processes with equally high-priority (Ramamritham et al., 1998). While using systems with multiple processors may provide some benefit, they do not alter the fundamental problem. System developers have no way to control the hardware outside of the capabilities provided by the running OS.

#### 2.1.1. Rex

Rex is a major solution in the world of Sensorimotor Research. Based on a QNX RTOS, it is capable of displaying events, interacting with DIO, AIO, with a sub-millisecond reliability. It was our main inspiration for this project and most of our objectif was to program a free alternative containing as many elements from Rex as possible, in an open, improvable, configurable framework.

#### 2.1.2. Matlab & PsychoToolBox

Matlab is a possible platform for such projects, nevertheless, there are notable limitations. Windows or Mac cannot support hard real-time operation. Therefore, while sub-millisecond jitter is acceptable in many, if not most, psychophysical settings, there are nonetheless many potential applications for which the software described here would not be suitable. In particular, experiments that must provide feedback within a very small temporal window (for example, to influence an on-going synaptic event) would find 2-5 ms jitter simply too variable. Furthermore, likewise, delivering feedback that requires a great deal of processing could potentially incur unacceptably long delays. There is a potential risk of blind period at the beginning of each behavioral tracking episode. Other limitations include the current inability to display movies or translating visual stimuli while simultaneously tracking behavioral signals. In addition, behavioral signals are not currently stored during the inter-trial interval. The ability to store analog signals continuously would benefit not only behavioral signals, but neurophysiological ones as well. In other words, although many acquisition cards are clearly capable in terms of number of channels, sampling rates and PC storage of recording neural data alongside behavioral signals, no support has been built-in for this purpose. Such a capability would likely be useful for many potential applications.

### 2.2. Our Solution: Rexeno

#### 2.2.1. General Presentation

We designed Rexeno’s structure to be flexible and compatible for many different types of experiments. One can plug about any analogical signal as an input (in Figure 1, we listed EEG, Eye Movements and Respiratory Movements, but these are just examples.) We also wanted to make the system compatible with any other hardware that requires triggering though digital signals (for example: Electrical stimulator devises, Transmagnetic stimulator, or Plexon hardware for single unit, local field potential or electroencepahlographical signals). We will show in subsection 4.2 how to configure these Strobes with Rexeno.

**Figure 1.**
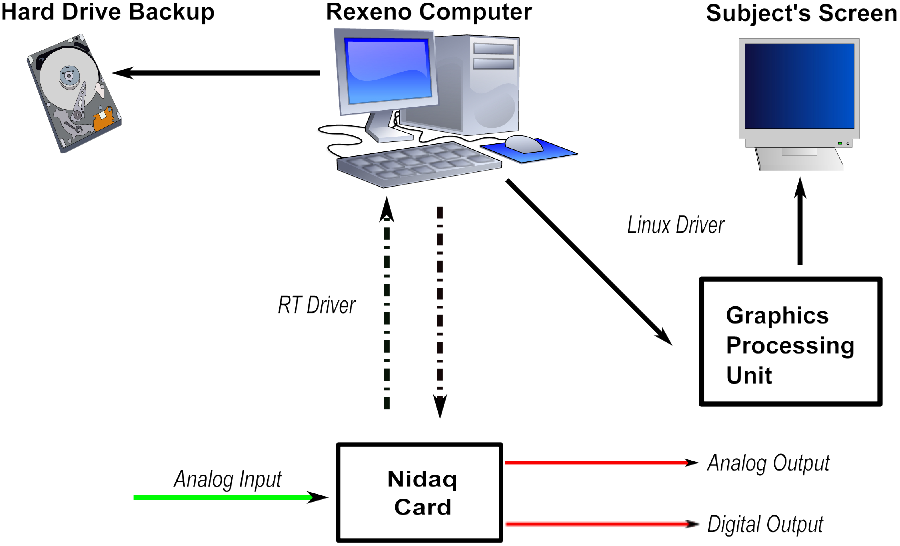
Rexeno Setup

#### 2.2.2. Hardware

Prioritizing portability, we interfaced high quality hardware that was already commonplace in the world of visual cognition study. Here is a list of the hardware used for our system:

- **Display**: CRT Screen
- **Data Acquisition**: National Instruments DAQ NI 6220
- **Graphic Card**: NVidia
- **CPU**: IntelXeonE5405

#### 2.2.3. Software & Solutions

Rexeno uses several freely available libraries, compatible with mainstream and high quality hardware, in order to obtain the best performances possible. Here are the main software ressources that we used in order to create our program:

- **Xenomai 2.6.0**: One of the most interesting options in Real-Time Operating Systems. Enables us to control software behabiour down to the μsec. Also provides real-time drivers for Analog/Digital interactions with AD cards.
- **OpenGL 2.0**: The main open-source graphical library. This 2.0 version allows us to display about anything and also provides powerful solutions to accelerate complicated rendering calculation with the graphic card using a dynamic pipeline (shader programming with GLSL)

#### 2.2.4. Verification methodology

We use this software on several different machines dedicated to neurophysiology in humans or non-human primates. This software has been very adept at the creation of basic sensori-motor tasks, and is especially useful for the creation of cognitive tasks with greater numbers of stimuli and contingencies. These tasks are often coded within an hour, and modifications are simple to test. Because different behavioral tasks can potentially place heavy demands on different aspects of the operating system and hardware (e.g., varying graphics, disk and memory use), end-users should not take observed timing accuracy in one task as direct evidence of satisfactory accuracy in another; thorough testing must be performed to assess the performance of new behavioral paradigms and new hardware configurations. The occurrence of temporal slips (unexpectedly increased latencies) often can be detected using time-stamps placed after critical behavioral events. These mark an event with reference to the deterministic system clock. A delay in the appearance of an expected time-stamp can then be used to reject trials in which timing constraints were not met. Of course, a delayed time-stamp could also represent a false-alarm when the delay occurred in the processing of that time-stamp itself and not in the preceding event. Fortunately, as we found above, such temporal slips can be made very infrequent, and are rarely longer than a millisecond. Once appropriate care has been taken to ensure accuracy in the three domains that are most likely introduce temporal jitter and error (video output, data sampling, and software), the reliance on a high-level language for behavioral control offers numerous benefits aside from simply the ease of task coding and portability across a wider range of hardware platforms. In particular, the simplicity with which new features can be coded encourages the development of new functions that improve usability and record keeping. While in principle such benefits could be realized in a low-level language, in practice, the difficulty and time-consuming nature of programming in such a language hinders their development by those who would like to spend their time designing and carrying out experiments rather than tweaking software.

## 3. Constraints

### 3.1. Definitions

As can be seen (in Figure 1), the interaction with the environment can be summerized with the following outputs

- **Display**: What appears on the subject’s screen.
- **Backup**: What was recorded by the machine
- **Digital Triggers**: Sent by the NIDAQ card.

Rexeno’s goal is to bring Hard Real-Time capability to these outputs.

Definition: A program is said to respect hard real time constraints when it’s time of execution is deterministic. The current system uses Xenomai’s Analogy Drivers for the interaction with NIDAQ card, so we know than these programs are real time compatible[?]. The interaction with the screens use NVidia proprietary drivers that do not offer real time guarentee (which is normal, this was not what they were designed for). Because of these drivers, it is not possible today to guarentee hard real time constraints while displaying something on a screen, what our tests will have to do is define under what conditions, jitters are still acceptable.

### 3.2. Testing the system

In order to verify the accuracy of our system, we designed the Double Flash Protocole, which consists of 2000 trials. Each trial consists of two successive flashes (duration = 50 frames ~ 833 ms). Time between front edge of the flash is 100 frames (≈ 1666.667ms). A photodiode is placed on the screen so that the flash creates a tension at the terminals of the photodiode. This tension was recorded at the same time by the Rexeno System, and by a Plexon Acquisition Unit. We can see a typical trial recorded on the Plexon unit in Figure 3a.

**Figure 2.**
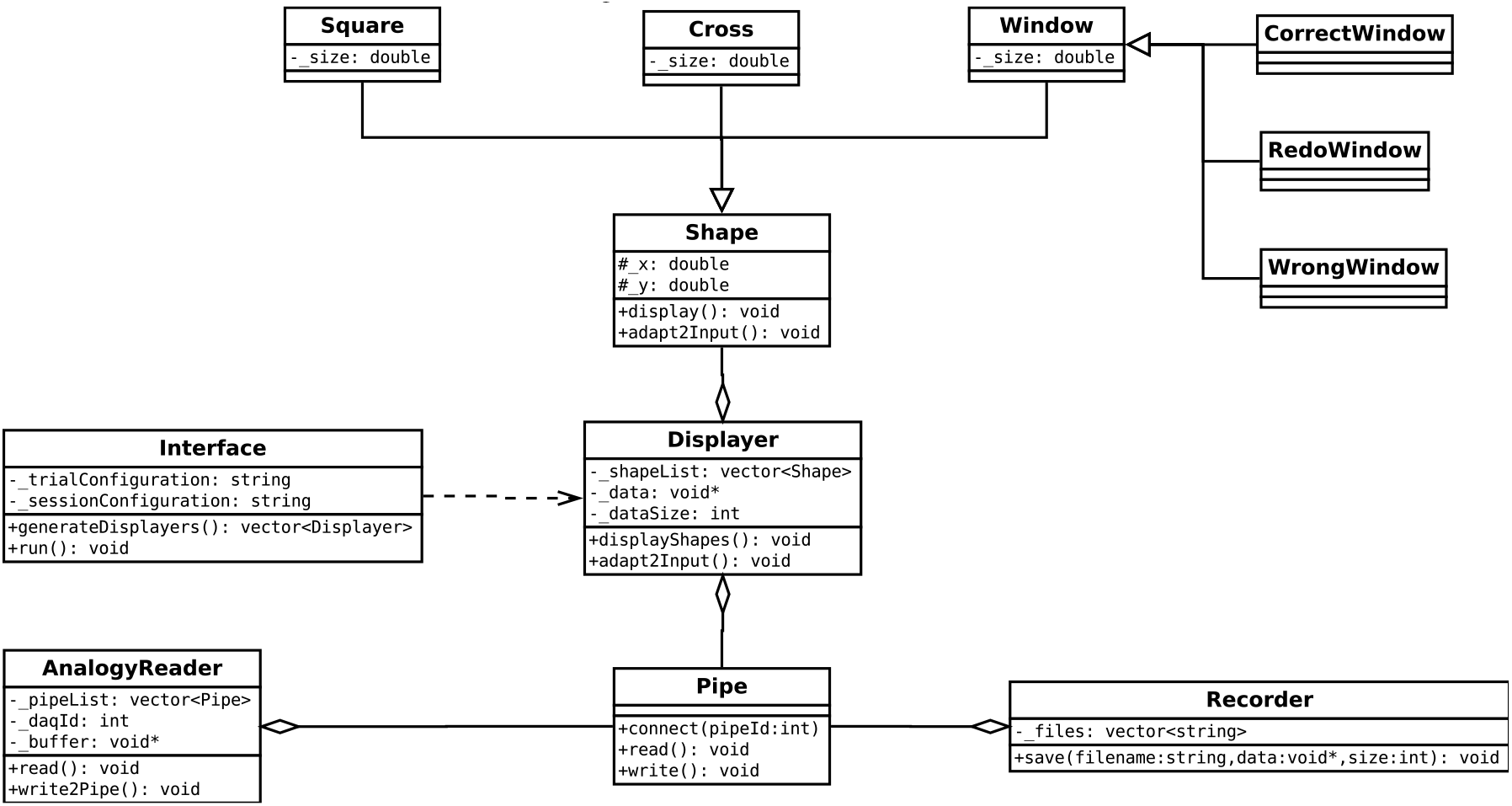
Software Architecture

**Figure 3.**
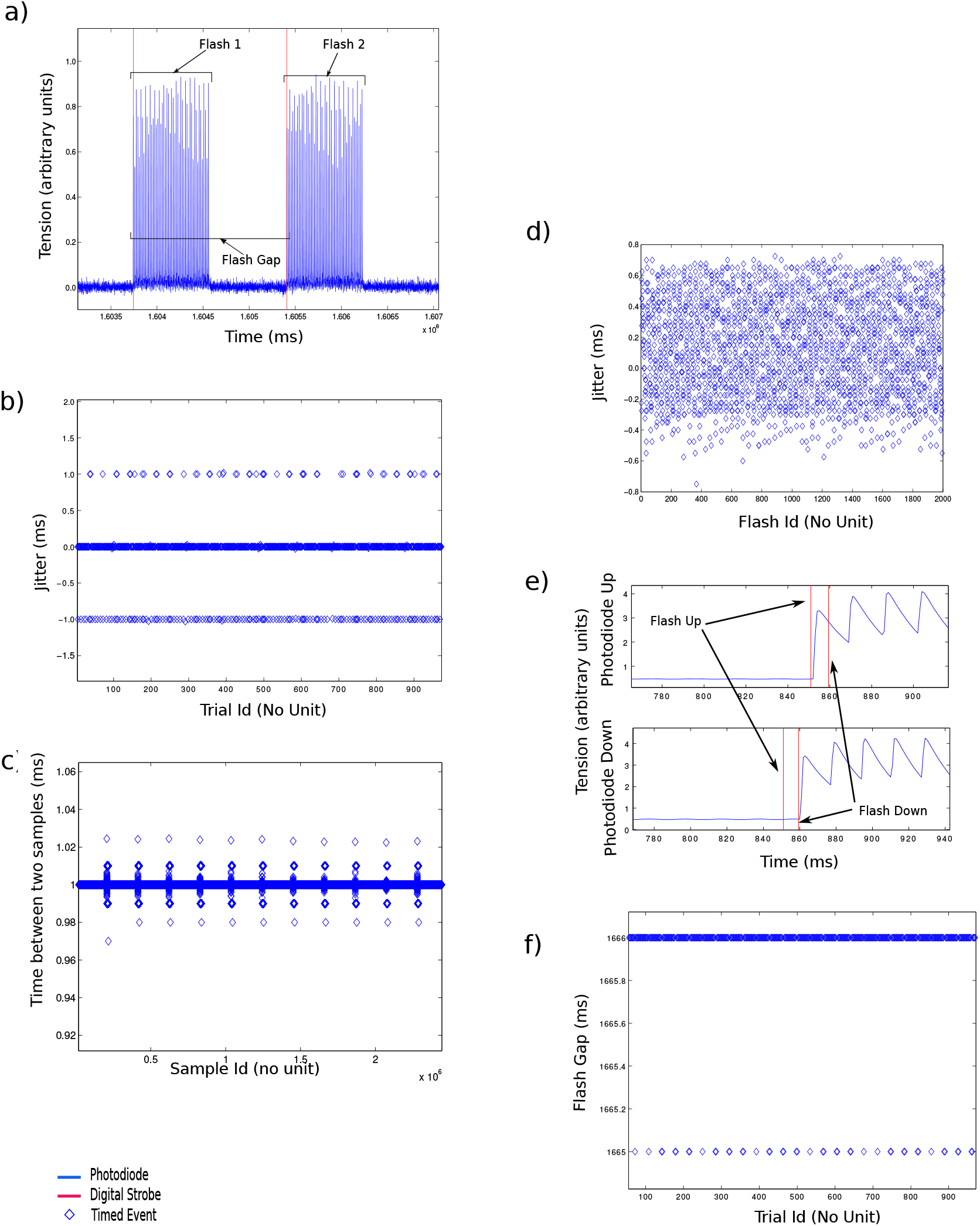
a,b,d,f - Two flashes at the same position separated by a time gap e - Two flashes on the same display frame but different positions c - Regular acquisition

It is designed to evaluate jitter between input and:

- Hard Drive Backup
- Displaying
- Digital output

### 3.3. Specific Hardware Interactions

#### 3.3.1. Hard Drive Backup Description

In a given trial, we wish to evaluate how precise the hard drive recording is. This task requires interaction with the Nidaq card and with the Hard Drive hardware. As said before, the Nidaq card has Real-Time compatible hardware (the Analogy drivers). This is not the case of the Hard Drive which cannot write in real time. This could be a problem because RT program is only as strong as it’s weakest link. A “seek” operation requires about 5ms (depending on the hardware configuration[?]), which is obviously not enough for 1000 Hz recording (writing on several files might create several “seek” operations on the hard drive at a given time, leading to data loss). Our solution was to use the IOstream C++ Library, which creates a buffer capable of saving on RAM memory large quantity of data and writing several Nidaq acquisitions to the disk using a single “write” operation. Another hardware specification indicates: “Host to/from drive (sustained) = 200Mb/sec“.

On the other hand, we aim for a 1000 Hz acquisition frequency (on 8 analogy channels) which creates the following quantity of data:

With:

- *N_Channels_* = 8
- *Size_Single_Acquisition_* = 8 Bytes
- *Frequency* = 1000Hz
- Θ represents the data’s encoding function^1^

Thanks to (3), we know we should be able to write all data in time without any loss (provided we use the streaming buffer technique). In the next two paragraphs, we will check that this was correctly implemented.

*Extracting data*

In order to check that the recorded manifestations have coherent timestamps, we decided to create an algorithm that will detect the Flash’s relative timestamps.

The result of this algorithm is shown in Figure 3b.

Our acquisition frequency being 1000 Hz (on the plexon and on the rexeno system,) the ±1 ms jitter is expected and compatible with Visual Cognition Studies (a visual saccade has a typical duration of 10ms).

#### 3.3.2. On-Screen Displaying

Displaying a stimuli is a task that requires interaction with a screen. This is done using the OpenGL2.0 library which will interact with the Nvidia graphic card’s drivers. The problem is the same as it was with the Hard Drive: the driver is not real-time compatible. In order to obtain deterministic displaying of stimuli, we took advantage of the CRT screen hardware which functions with a 60Hz displaying frequency. Our technique was simple and exploited the OpenGL’s double buffer capacity: if the protocol needs to draw a stimuli at the *n^th^* frame, we wait until the (n ‒ 1)*^th^* frame and draw the corresponding stimuli on the back buffer which will automatically be displayed on the CRT screen at the next frame.

The flash gap was supposed to be 100 frames at 60Hz, On Figure 3f is the evaluation of this gap using the Plexon recording and the FrontEdge algorithm. The 1000Hz acquisition frequency creating again a ± 1 ms jitter.

#### 3.3.3. Digital Output

Triggering digital output uses only the Analogy RT drivers. So we can control time very precisely. In order to trigger events with displayed stimuli, we wait for a vBlank synchronisation to occur, wait a certain amount of time depending of the CRT’s frequency and send the digital pulse. To evaluate the jitter, we recorded the strobe events on the plexon and compared these timestamps to the ones returned by our FrontEdges function. The result of this difference can be seen on Figure 3b.

#### 3.3.4. Vertical Synchronisation & Tearing

In order to fully control what is displayed on the subject’s screen, we had to use the VBlank synchronisation of the NVidia drivers. If enabled, the graphic card waits for the VBlank event (cathod ray beam is turned off for repositioning at the top left of the screen) before starting to draw anything. This avoids the tearing effect. We used an experiment where two flashes (one on the “up” half of the screen, the other on the “bottom” side) were supposed to appear on screen at the same frame. Theoretically, the “up” flash should appear before the “down” flash because of the VBlank synchronisation. With a photodiode placed at various positions, we decided to verify that assertion. Results are on Figure 3e.

### 3.4. Conclusion

Even if it is impossible today to control in Hard Real Time every feature of the Rexeno station, we can emulate real time by exploiting various techniques on a regular x86 computer.

## 4. How to use Rexeno

### 4.1. Using Rexeno

As we have seen, creating a fully reliable real time environment can be a challenging task. That is why we have decided to fully package all these algorithms into one piece of software and distribute it freely to anyone who would be interested. Making it, we hope, a good choice for people interested in setting up an experiment where precise timing is necessary.

### 4.2. Defining a task

The user can define his protocole though a GUI that creates configurations files. These will be in charge of describing the different trials.

**Figure 4.**
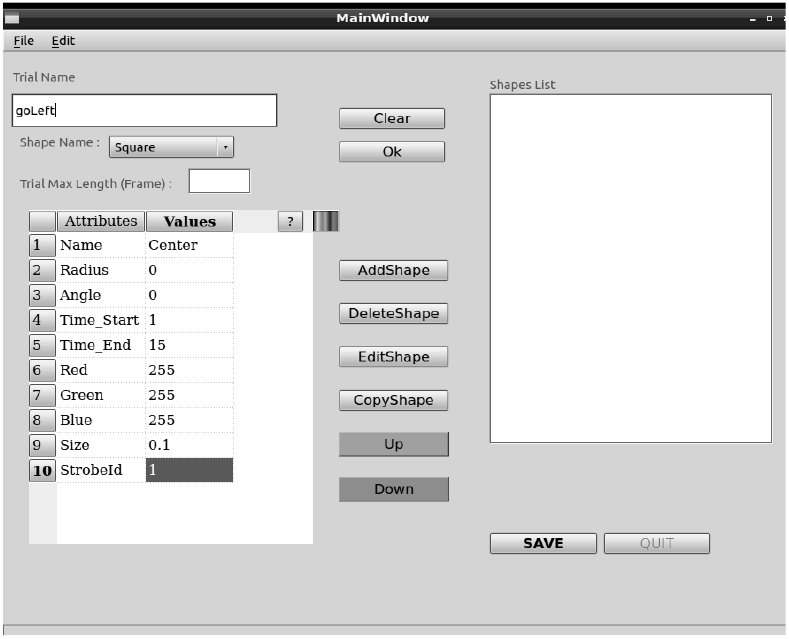
Interface for trial creation

Here is an example of a configuration file created by the interface:

idtrial1 300 GO_TARGET1

id1 Square Target 0.97 0 End_Fixation End_Target 0 255 0 0.03 3

id2 Cross Eye 0 0 0 10000 255 0 0 0.1

id3 FixationWindow Fixation 0 0 0 End_Fixation 255 255 0 0.32 0.32 Fixation_Duration

id4 NeutralWindow Neutral 0 0 0 End_Fixation 0 200 0 0.32 0.32

id5 RedoWindow Redo 0 0 100 End_Fixation 200 0 0 2 2 0

id6 Square Fixation 0 0 0 End_Fixation 0 255 0 0.03 1

id7 CorrectWindow CorrectWindow 0.97 0 End_Fixation End_Target 255 255 0 0.5 0.5 Target_Time

id8 Time 300

id9 Variable GO_TARGET1 1 endtrial

With such a file, we can launch the task with a subject. The main events are presented in Figure ??

**Figure 5.**
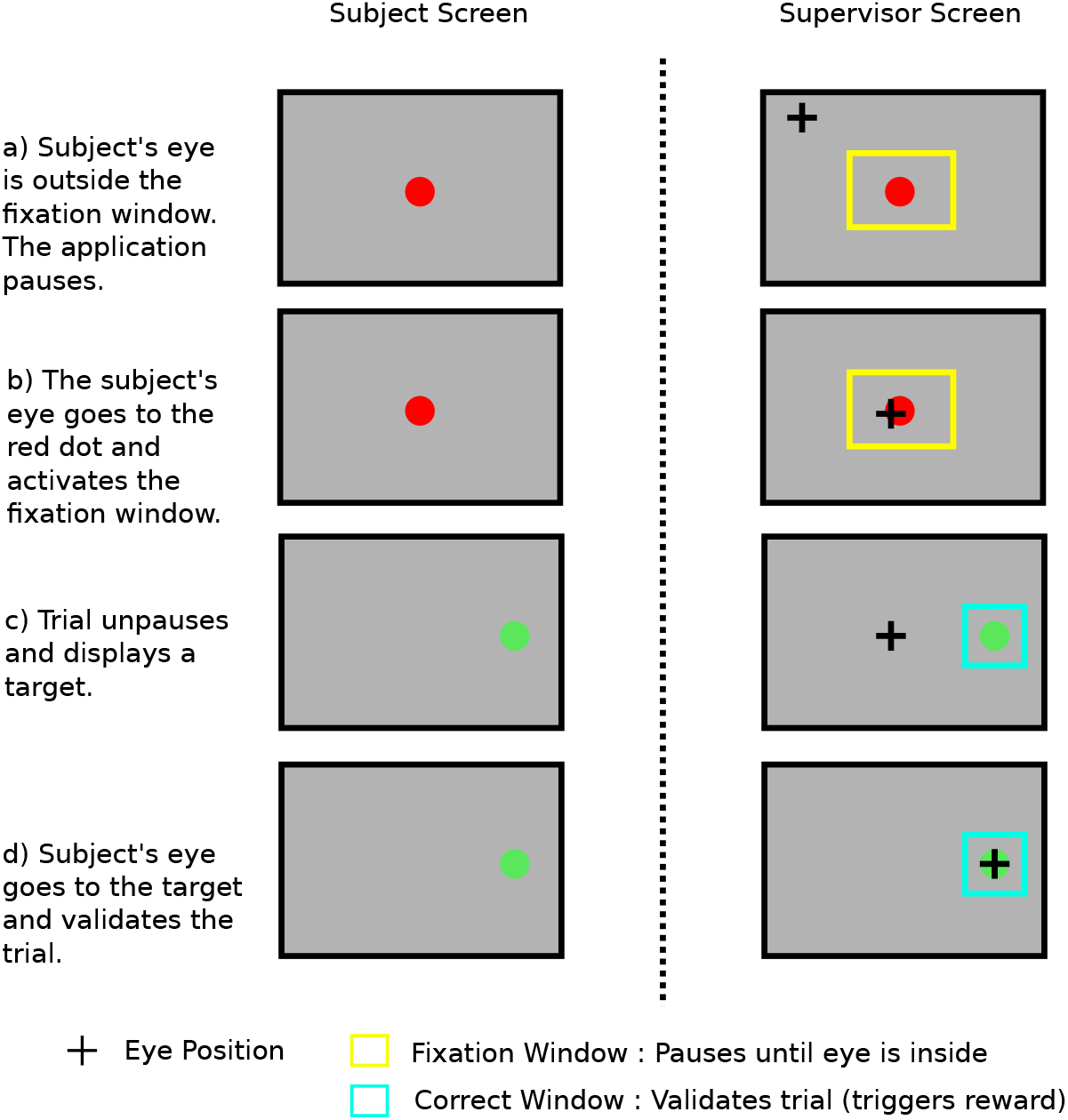
Interface for trial creation

## Footnotes

1 Depends on the entropy but it’s asymptot should be a constant

